# Subcellular dynamics and functional activity of the cleaved Na^+^ channel β1 subunit intracellular domain

**DOI:** 10.1101/2021.12.29.474414

**Authors:** Alexander S. Haworth, Samantha L. Hodges, Alina L. Capatina, Lori L. Isom, Christoph G. Baumann, William J. Brackenbury

## Abstract

The voltage-gated Na^+^ channel β1 subunit, encoded by *SCN1B*, regulates cell surface expression and gating of α subunits, and participates in cell adhesion. β1 is cleaved by α/β and γ-secretases, releasing an extracellular domain and intracellular domain (ICD) respectively. Abnormal *SCN1B* expression/function is linked to pathologies including epilepsy, cardiac arrhythmia, and cancer. In this study, we sought to determine the effect of secretase cleavage on β1 function in breast cancer cells. Using a series of GFP-tagged β1 constructs, we show that β1-GFP is mainly retained intracellularly, particularly in the endoplasmic reticulum and endolysosomal pathway, and accumulates in the nucleus. Reduction in endosomal β1-GFP levels occurred following γ-secretase inhibition, implicating endosomes, and/or the preceding plasma membrane, as important sites for secretase processing. Using live-cell imaging, we report β1ICD-GFP accumulation in the nucleus. Furthermore, β1-GFP and β1ICD-GFP both increased Na^+^ current, whereas β1STOP-GFP, which lacks the ICD, did not, thus highlighting that the β1-ICD was necessary and sufficient to increase Na^+^ current measured at the plasma membrane. Importantly, although the endogenous Na^+^ current expressed in MDA-MB-231 cells is TTX-resistant (carried by Na_v_1.5), the Na^+^ current increased by β1-GFP or β1ICD-GFP was TTX-sensitive. In addition, β1-GFP increased mRNA levels of the TTX-sensitive α subunits *SCN1A*/Na_v_1.1 and *SCN9A*/Na_v_1.7. Taken together, this work suggests that the β1-ICD is a critical regulator of α subunit function in cancer cells. Our data further highlight that γ-secretase may play a key role in regulating β1 function in breast cancer.

## Introduction

Voltage-gated Na^+^ channels (VGSCs) are heteromeric complexes consisting of Na^+^-conducting α subunits (Na_v_1.1-1.5, encoded by *SCN1A-5A*, and Na_v_1.6-1.9, encoded by *SCN8A-11A*) and non-pore-forming β subunits (β1-β4, encoded by *SCN1B-4B*) (1). The inward Na^+^ current carried by VGSCs is responsible for membrane depolarisation during action potential initiation (2). With the exception of the β1 alternative splice variant β1B, β subunits are single-pass transmembrane proteins with a large, extracellular immunoglobulin (Ig) domain and are thus members of the Ig superfamily of cell adhesion molecules (CAMs) (3). The β subunits regulate α subunit trafficking (4-6), cell type-specific gating and kinetics (7-15), mechanosensitivity (16,17), and glycosylation (18). In addition to regulating α subunit function, β subunits also function as CAMs, regulating cell-cell and cell-matrix adhesion via interaction with an array of other CAMs, neurite outgrowth, neuronal pathfinding, fasciculation, and cell migration (6,15,19-25). β1-mediated cell adhesion interactions also recruit ankyrin to adhesion contacts and promote neurite outgrowth via activation of fyn kinase (21,26-28).

Variants in genes encoding VGSC α and β subunits occur in various excitability-linked disorders, including epilepsy and cardiac arrhythmia (29,30). Variants in *SCN1B*, encoding β1, are associated with developmental and epileptic encephalopathy (DEE56), early infantile DEE (EI-DEE), genetic epilepsy with febrile seizures plus, atrial fibrillation and Brugada syndrome (31-34). *Scn1b* null mice display EI-DEE and disrupted neuronal pathfinding and fasciculation, as well as altered cardiac excitability (12,13,35,36). VGSCs are also aberrantly expressed in cancer cells (37). β1 is upregulated in breast cancer tissue compared to healthy tissue (38). β1 overexpression in metastatic MDA-MB-231 breast cancer cells increases Na^+^ current, without altering gating kinetics, induces outgrowth of neurite-like processes *in vitro* and increases tumour growth and metastasis *in vivo* (38,39). Taken together, these observations highlight that *SCN1B* plays a key role in regulating pathophysiological behaviour of excitable and non-excitable cells.

β1 interacts with α subunits via extracellular and intracellular sites (4,40,41). Extracellularly, β1 contacts extracellular loops within DI, DIII, DIV of Na_v_1.4, as well as within the DIII transmembrane domain (42,43). Although an intracellular α-β1 interaction site has yet to be resolved, a soluble polypeptide representing the intracellular C-terminus of Na_v_1.1 co-immunoprecipitates with β1 (41). Furthermore, deletion of the β1 intracellular domain (ICD) attenuates β1-Na_v_1.2 interaction (4). The locations of α-β1 extracellular/intracellular interaction sites are intriguing because β subunits are substrates for regulated intramembrane proteolysis by sequential activity of α- or β-secretase then γ-secretase, releasing the extracellular Ig domain and then the soluble ICD, respectively (44-46). In addition, palmitoylation of β1 is required for its proteolytic processing at the plasma membrane (47). The soluble extracellular β1 Ig domain has been shown to promote neurite outgrowth (3,24). The β1-ICD and β2-ICD have been shown to accumulate in the nucleus, regulating transcription (46,48). Thus, proteolytic processing of β1 plays a key role in regulating adhesion, neurite outgrowth and gene transcription.

γ-secretase activity promotes cancer progression via activation of Notch signalling, and several γ-secretase inhibitors have been pursued in clinical trials (49). Moreover, aberrant β1ICD-mediated transcriptional changes may promote β1-linked pathologies, including epilepsy, cardiac arrhythmia, and cancer (46). Here, we sought to determine the effect of secretase cleavage on β1 function in breast cancer cells. Using a series of GFP-tagged β1 constructs stably expressed in MDA-MB-231 cells, we found that full-length β1-GFP is mainly retained intracellularly, particularly in the endoplasmic reticulum and endolysosomal pathway, and accumulates in the nucleus. Pharmacological inhibition of γ-secretase cleavage decreased β1ICD-GFP levels but had no effect on spatiotemporal cycling dynamics of β1-GFP and did not alter Na^+^ current. Using live-cell imaging, we report specific β1ICD-GFP accumulation in the nucleus. Furthermore, β1-GFP or β1ICD-GFP overexpression increased Na^+^ current, whereas β1STOP-GFP, which lacks the ICD, did not, thus highlighting a requirement for the ICD to promote VGSCs at the plasma membrane. Importantly, although the endogenous Na^+^ current expressed in MDA-MB-231 cells is tetrodotoxin (TTX)-resistant (carried by Na_v_1.5) (50,51), the Na^+^ current increased by β1-GFP or β1ICD-GFP was TTX-sensitive. Taken together, this work suggests that the proteolytically cleaved β1-ICD is a critical regulator of α subunit function in breast cancer cells.

## Results

### Plasma membrane expression and activity of β1-GFP

In this study, we used over-expression of β1-GFP in the MDA-MB-231 cell line as a model system in which to study functional consequences of proteolytic processing of β1 by secretase cleavage. MDA-MB-231 cells provide a unique model system to analyse β1 function for two reasons. Firstly, there is low endogenous β subunit expression in this cell line, thus enabling introduction of engineered β1 constructs (39). Secondly, endogenous expression of functional α subunits in MDA-MB-231 cells negates the requirement for exogenous α subunit expression and ensures a native trafficking pathway is present for α subunits to reach the plasma membrane. Initially, Na^+^ currents in MDA-MB-231-β1-GFP cells were compared to control MDA-MB-231-GFP cells using whole cell patch clamp recording. Peak Na^+^ current density in MDA-MB-231-β1-GFP cells was 3-fold greater than cells expressing GFP alone, -16.80 ± 8.20 pA/pF vs. -5.16 ± 2.01 pA/pF (P < 0.01; n = 8; t-test; Figure 1A-C). These data suggest that β1-GFP increases α subunit expression at the plasma membrane, in agreement with previous observations (39). β1-GFP over-expression did not affect the voltage at activation, voltage at half-maximal activation, rate of activation, voltage at half-inactivation, rate of inactivation, time to current peak, or membrane capacitance (Figure 1D-G and I-M). However, β1-GFP over-expression caused a hyperpolarisation of the voltage at Na^+^ current peak (P < 0.05; n = 8; t-test; Figure 1H), although the small shift, together with the lack of change in voltage-dependence of activation, suggests that this change is unlikely to be physiologically important. β1-GFP over-expression also accelerated recovery from inactivation (P < 0.01; n = 8; t-test; Figure 1N, O).

**Figure 1.**
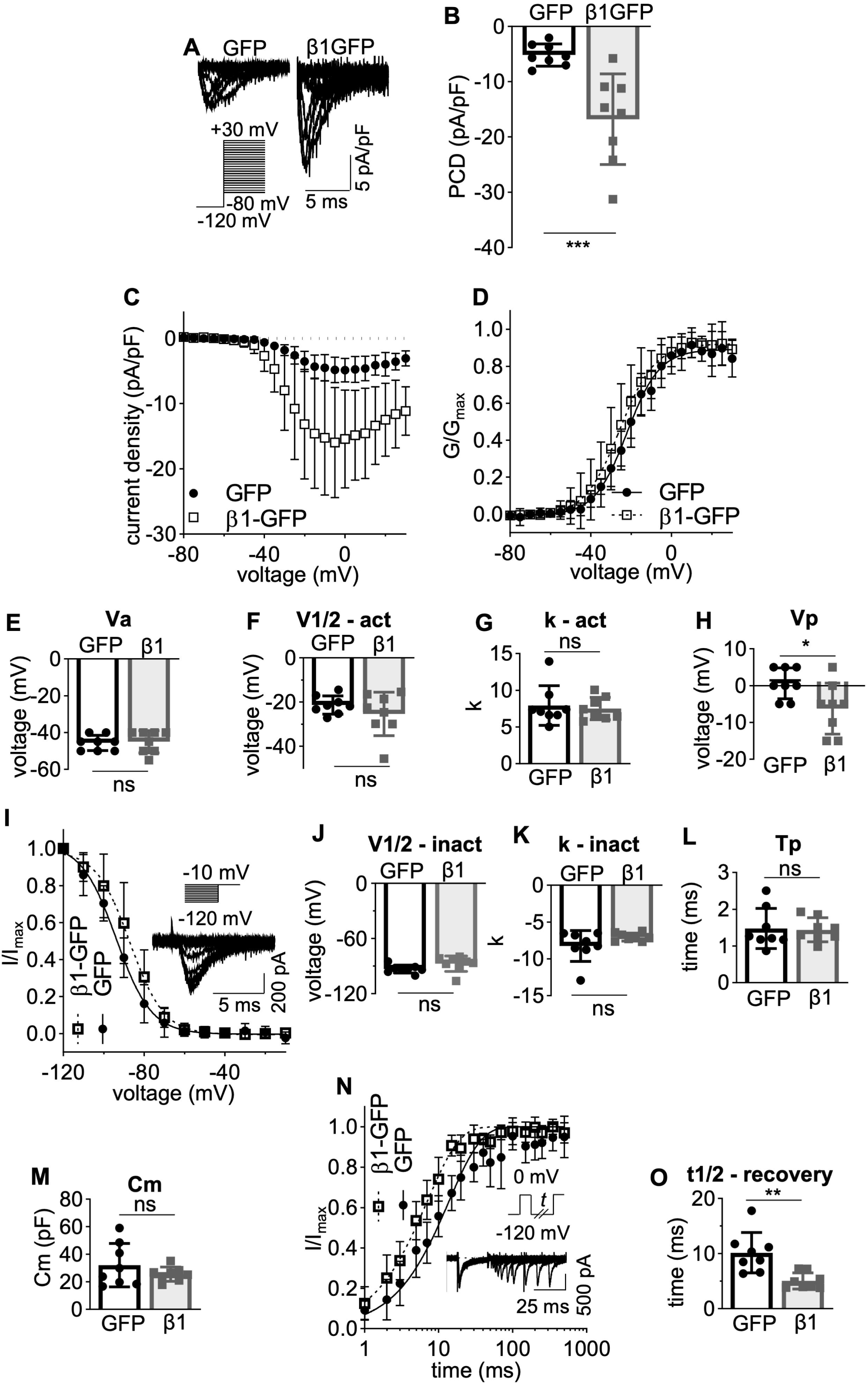
Electrophysiological properties of MDA-MB-231-β1-GFP cells. (A) Representative whole-cell Na^+^ currents in MDA-MB-231-GFP and MDA-MB-231-β1-GFP cells, following depolarisation between -80 mV and +30 mV for 250 ms from -120 mV. (B) Peak current density (PCD). (C) Current (I)-voltage (V) relationship. (D) Conductance (G)-voltage (V) relationship. (E) voltage at activation (Va). (F) voltage at half-maximal activation (V1/2-act). (G) Slope factor of activation (k – act). (H) Voltage at peak current (Vp). (I) Steady-state inactivation. Cells were depolarised at -10 mV following a 250 ms holding voltage of between -80 mV and +30 mV. (J) Voltage at half-maximal inactivation (V1/2-inact). (K) Slope factor of inactivation (k – inact). (L) Time to peak at 0 mV (Tp). (M) Whole cell capacitance (Cm). (N) Recovery from inactivation. Cells were depolarised to 0 mV for 25 ms, then held at -120 mV for *t* s before second depolarisation to 0 mV. *t* ranged from 1-500 ms. (O) time taken for half-maximal recovery. Data are presented as mean ± SD (n = 8, N = 3). Activation and inactivation curves are fitted with a Boltzmann function. ns = not significant, * = P < 0.05, ** = P < 0.01, unpaired t-test.

The observations that β1-GFP over-expression (i) increases Na^+^ current and (ii) promotes transcellular adhesion of MDA-MB-231 cells (39) suggest that it is functionally active at the plasma membrane in this cell line. We therefore examined the subcellular localisation of β1-GFP, initially focusing on plasma membrane expression. Surprisingly, when live MDA-MB-231-β1-GFP cells were stained with the lipid dye FM4-64, no overlap in fluorescence was detected at the plasma membrane, whereas robust co-localisation was observed within internal vesicles (Figure 2A). In fact, line profile analysis revealed that peak plasma membrane FM4-64 fluorescence and β1-GFP fluorescence were offset by ∼500 nm (Figure 2B), suggesting that β1-GFP is not highly expressed at the plasma membrane relative to the cytosol. To ensure that the lack of surface β1-GFP abundance was not due to FM4-64 quenching GFP fluorescence via FRET, FM4-64 was photobleached and the resulting change in GFP fluorescence monitored. Photobleaching of FM4-64 within internal vesicles caused a modest, but significant, 8.9 % increase in GFP signal (P < 0.05; n = 4; t-test; Figure 2C), suggesting that some FRET did occur between GFP and FM4-64. However, when FM4-64 was photobleached at the plasma membrane, no increase in GFP signal was detected (Figure 2C), ruling out GFP quenching by FM4-64 as an explanation for the low abundance of β1-GFP at the cell surface. In summary, although β1-GFP promotes Na^+^ current, most of this protein appears to be retained intracellularly in MDA-MB-231 cells. This observation agrees with a previous study in Madin-Darby canine kidney cells, which showed that β1 was retained intracellularly, unlike β2, which was enriched at the plasma membrane (52).

**Figure 2.**
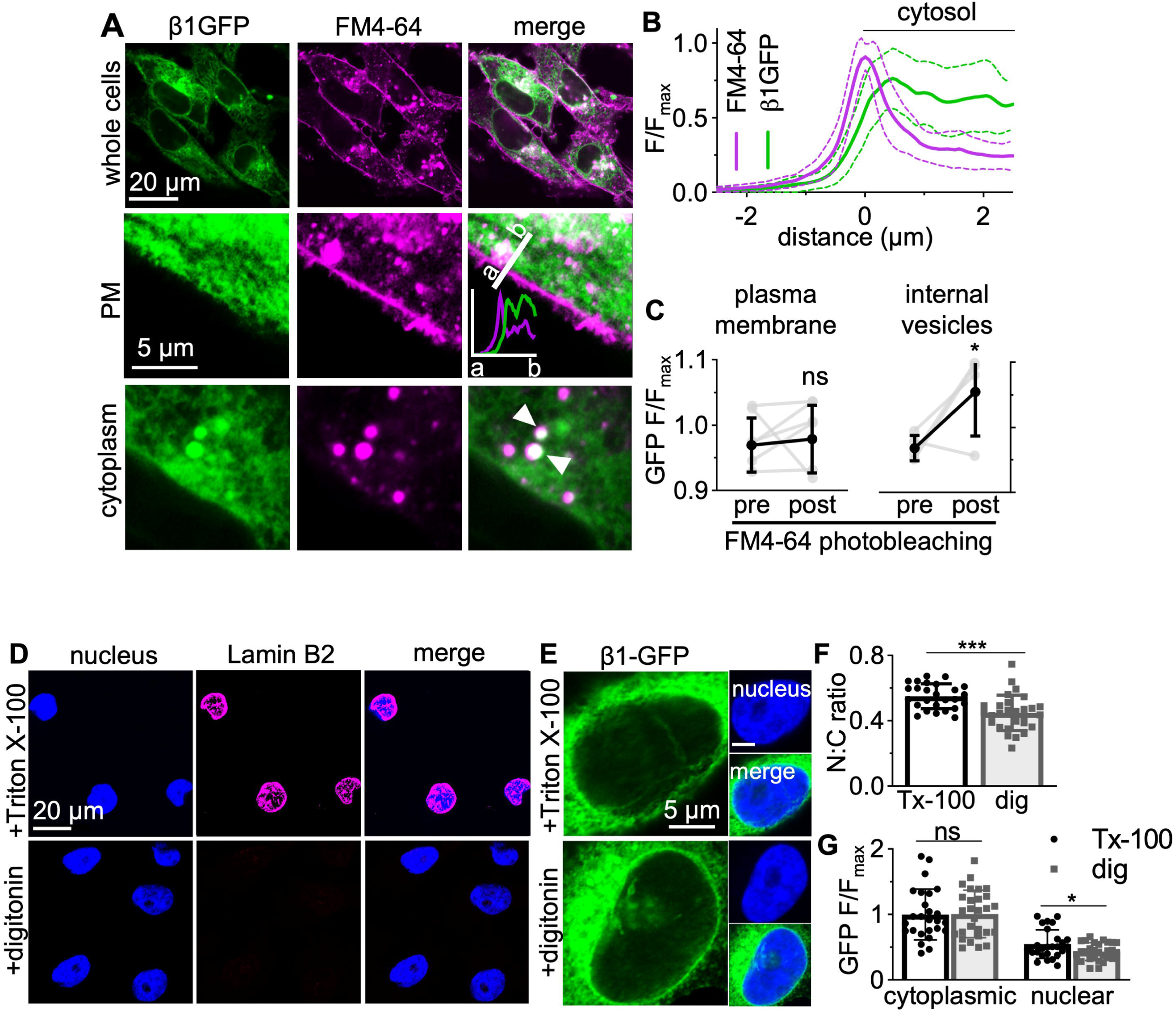
Plasma membrane and nuclear localisation of β1-GFP. (A) Confocal images of live MDA-MB-231-β1-GFP cells stained just prior to imaging with FM4-64 (120 nM). Panels include images of whole cells, the plasma membrane (PM) and the cytoplasm. Line profile shown on plasma membrane image demonstrates GFP/FM4-64 fluorescence. White arrowheads denote areas of β1-GFP/FM4-64 overlap. (B) A 5 μm line profile across the plasma membrane of live MDA-MB-231-β1-GFP cells. Data are mean (solid line) ± SD (dotted line), n = 10 cells. (C) Quantification of FRET between GFP and FM4-64 within intracellular vesicles and at the plasma membrane. Regions of interest were photobleached using 100 iterations of a 561 nm laser (100 % laser power) to achieve 80-90 % bleaching of FM4-64. GFP fluorescence (normalised to fluorescence of the first frame) between the pre-bleach and post-bleach recording were compared using an unpaired t-test. n = 5–6. Data are mean ± SD. (D,E) MDA-MB-231-β1-GFP cells were fixed and permeabilised with Triton X-100 (0.3 %) or digitonin (50 μg/ml). Cells were labelled with antibodies for Lamin B2 (magenta) or GFP (green). (F) Quantification of the nuclear:cytoplasmic (N:C) signal density ratio between Triton X-100 and digitonin-permeabilised cells. Data are mean ± SD (n = 27 – 28, N = 3). (G) Comparison of the cytoplasmic fluorescence intensity and nuclear fluorescence intensity between MDA-MB-231-β1-GFP cells permeabilised with Triton X-100 or digitonin. Data are mean ± SD (n = 27 – 28, N = 3). ns = not significant, * = P < 0.05, *** = P<0.001, unpaired t test.

### Subcellular distribution of β1-GFP

The β1-ICD and β2-ICD secretase cleavage products localise to the nucleus of heterologous cells and alter gene transcription (46,48). We therefore next investigated whether any β1-GFP signal localised to the nucleus in MDA-MB-231 cells. Prior to anti-GFP antibody labelling, cells were permeabilised with either Triton X-100, which permeabilises all cellular membranes, permitting access to nuclear antigens, or digitonin, which does not permeabilise the nuclear membrane, preventing access to nuclear antigens (53). The inner nuclear membrane protein, lamin B2, was labelled strongly in Triton X-100 permeabilised cells, but not digitonin-permeabilised cells, confirming that digitonin restricted antibody access to nuclear antigens (Figure 2D). Labelling with the anti-GFP antibody revealed a small but statistically significant 18 % reduction in nuclear:cytoplasmic fluorescence intensity ratio in digitonin-permeabilised cells compared to Triton X-100 permeabilised cells (P < 0.001; n = 26-28; t-test; Figure 2E, F), suggesting that a fraction of β1-GFP is indeed present in the nucleus. Furthermore, cytoplasmic GFP fluorescence intensity was similar between the two permeabilization conditions (P = 0.86; n = 27; t-test; Figure 2G), whereas nuclear GFP fluorescence intensity was significantly reduced in digitonin-permeabilised cells by 23 % (P <0.05; n = 27; t-test; Figure 2G). Together, these data support the notion that there is a fraction of β1-GFP signal which localises to the nucleus in MDA-MB-231-β1-GFP cells.

We next studied co-localisation with specific organelle markers to further characterise the subcellular distribution of β1-GFP. Given that β1-GFP is a transmembrane protein, and an intracellular, punctate-like distribution is prominent in MDA-MB-231-β1-GFP cells (Figure 2A), we hypothesised that β1-GFP would be present within the endocytic pathway. Indeed, β1-GFP partially colocalised with the early endosome marker EEA1 (Figure 3A, F), suggesting that it is present within early endosomes following internalisation. Similarly, β1-GFP partially colocalised with the lysosomal marker LAMP1 (Figure 3B, F), suggesting that it also progresses to lysosomes for degradation. To further confirm lysosomal expression of β1-GFP, MDA-MB-231-β1-GFP cells were treated with chloroquine, an inhibitor of lysosomal degradation. Chloroquine treatment caused a characteristic swelling and vacuolization of lysosomes (54), resulting in an accumulation of β1-GFP within enlarged intracellular vesicles (Figure 3G). Cells were next labelled for marker proteins of the endoplasmic reticulum (calnexin), and *cis*- and *trans*-Golgi networks (GM130 and TGN46, respectively). β1-GFP partially colocalised with GM130 (Figure 3C, F); however, there was a more robust overlap between β1-GFP and TGN46 (Figure 3D, F) and calnexin (Figure 3E, F), suggesting that β1-GFP is more abundant in the *trans*-Golgi network and endoplasmic reticulum than the *cis-*Golgi.

**Figure 3.**
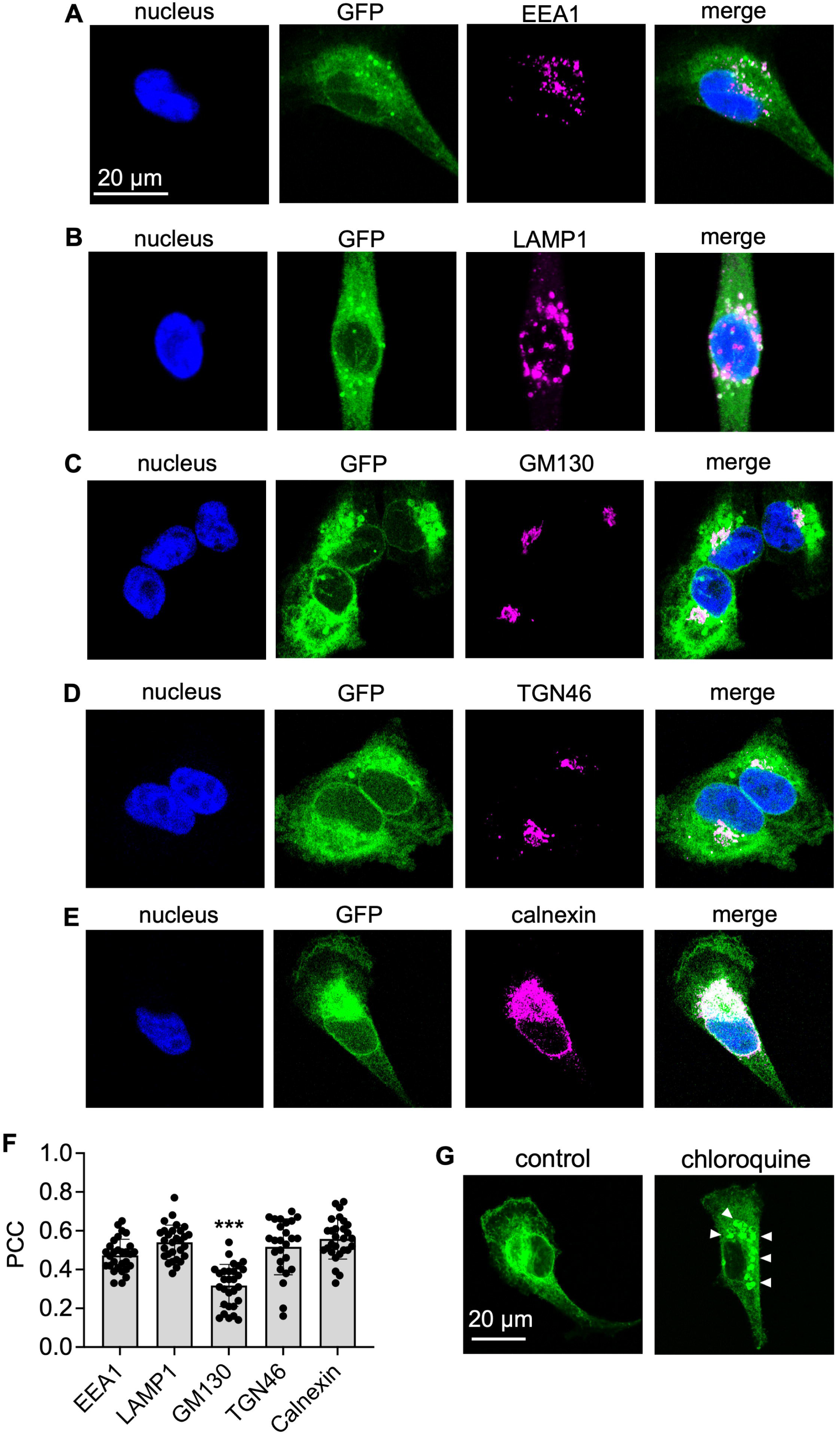
Subcellular distribution of β1-GFP. MDA-MB-231-β1-GFP cells were stained for (A) EEA1, (B) LAMP1, (C) GM130, (D) TGN46, (E) calnexin (all magenta), and DNA (DAPI, blue). Endogenous GFP is shown in green. Confocal microscopy with Airyscan technology was used for acquisition. (F) Quantification of the co-localisation of GFP and the indicated markers was calculated using Pearson’s correlation coefficient. Data are mean ± SD (n = 25 - 29, N = 3). * = P < 0.05; one-way ANOVA with Tukey’s test. (G) MDA-MB-231-β1-GFP cells were pre-treated with/without chloroquine (10 μM, 24h), then fixed and endogenous GFP detected using confocal microscopy with Airyscan technology. White arrowheads indicate enlarged lysosomes.

### Effect of γ-secretase inhibition on β1-GFP processing and function

Secretase cleavage remains an uncertain point of regulation for β1, and it is not fully understood whether secretase cleavage influences β1-mediated α subunit regulation. An α-subunit interaction site within the β1-ICD, which is responsible for α-subunit surface trafficking, presents the possibility of secretase-mediated regulation of Na^+^ current via β1 (4). In addition, γ-secretase inhibition prevents β2-mediated cell adhesion (45), and reduces β1-mediated neurite outgrowth (3), suggesting that the CAM function of β1 could also be regulated by secretase processing. To investigate these possibilities, we treated MDA-MB-231-β1-GFP cells with the γ-secretase inhibitor, DAPT (Figure 4A). DAPT treatment reduced the amount of β1ICD-GFP cleavage product present, increasing the C-terminal fragment (CTF):ICD expression ratio by 15-fold (P < 0.01; n = 4–5; t-test; Figure 4B). DAPT had no effect on the α-tubulin level (P =0.57; n = 4-5; t-test; Figure 4B), suggesting that the treatment did not alter total protein levels. We next tested the effect of γ-secretase inhibition on Na^+^ current in MDA-MB-231-β1-GFP cells. Peak current density of MDA-MB-231-β1-GFP cells treated with DAPT (-30.74 ± 10.16 pA/pF; n = 10) was no different to that of DMSO vehicle-treated cells (-26.89 ± 6.44 pA/pF; P = 0.29; n = 12; t-test; Figure 4C). This result suggests that γ-secretase activity is not involved in the β1-GFP-mediated increase in Na^+^ current. We also tested the effect of two other γ-secretase inhibitors, L-685,458 and avagacestat. Both compounds reduced the amount of β1ICD-GFP and increased the CTF:ICD expression ratio (P < 0.05; n = 3; one-way ANOVA; Figure 4D). Similar to DAPT, avagacestat treatment had no effect on peak current density (P = 0.11; n = 7-8; t-test; Figure 4E). In contrast, L-685,458 (10 µM) did inhibit peak current density in MDA-MB-231-β1-GFP cells (P < 0.01; n = 8-18; one-way ANOVA; Figure 4F, top sub-panel); however, it also inhibited peak current density in control MDA-MB-231-GFP cells (P < 0.01; n = 8-18; one-way ANOVA; Figure 4F, bottom sub-panel), suggesting the inhibition was independent of β1-GFP. At a lower dose (1 μM), L-685,458 did not inhibit peak current density in either MDA-MB-231-β1-GFP (P = 0.69; n = 8-18; one-way ANOVA; Figure 4F, top sub-panel) or MDA-MB-231-GFP cells (P = 0.16; n = 8-18; one-way ANOVA; Figure 4F, bottom sub-panel). None of the γ-secretase inhibitors affected channel recovery from inactivation (DAPT: P = 0.25, n = 8, t-test; avagacestat: P = 0.29, n = 5-8, t-test; L685,458: P = 0.11, n = 8, t-test; Figure 4G). In summary, pharmacological inhibition of γ-secretase caused a reduction in the level of the β1-ICD cleavage product; however, it had no detectable effect on Na^+^ current density or recovery from inactivation.

**Figure 4.**
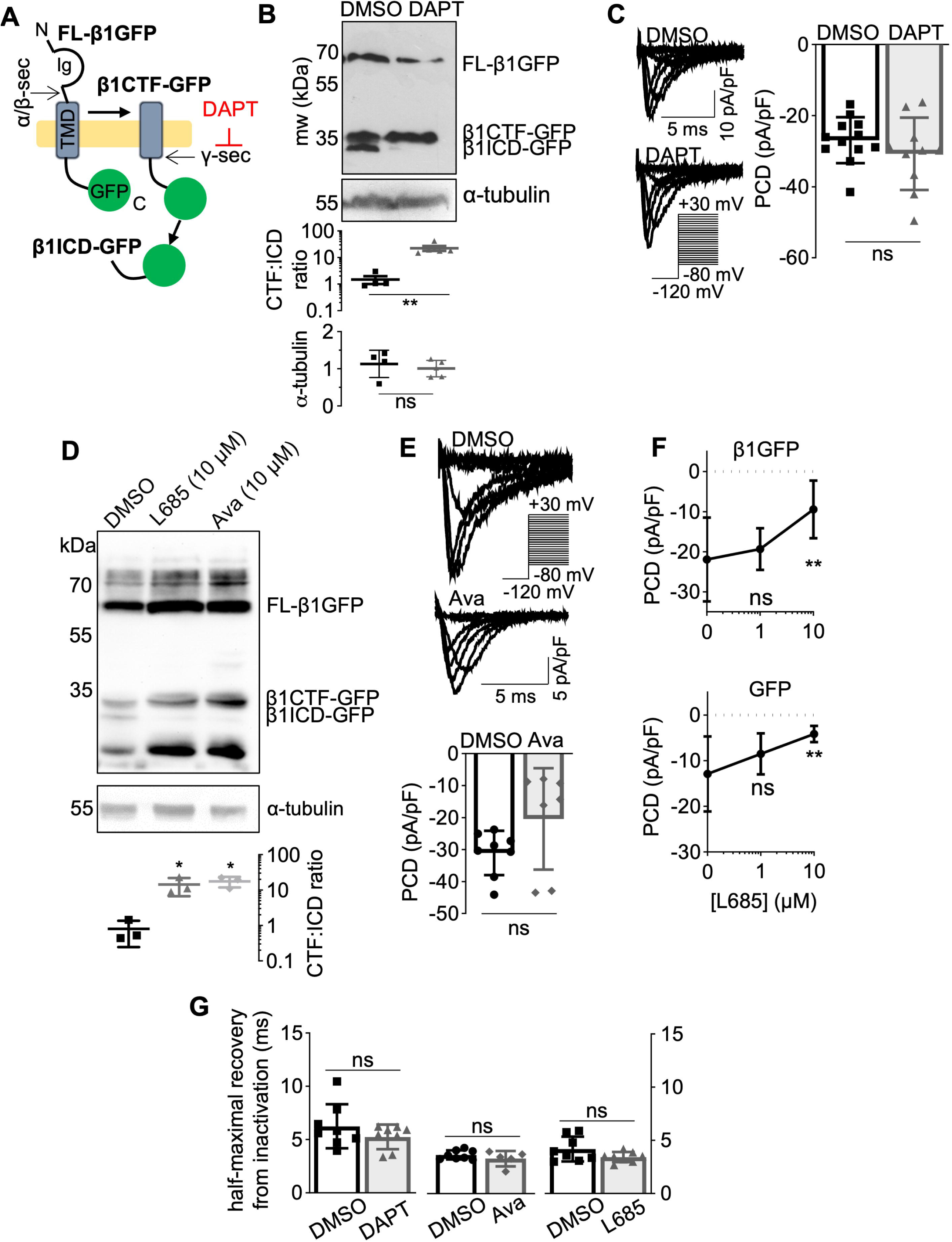
Effect of γ-secretase inhibition on β1-GFP function. (A) Schematic depicting sequential secretase processing of β1-GFP. FL: full length, CTF: C-terminal fragment, ICD: intracellular domain. Ig: immunoglobulin loop. γ-secretase is inhibited by DAPT. (B) Western blot of lysates from MDA-MB-231-β1-GFP cells treated with DMSO (0.01 % (v/v), 24 h) or DAPT (1 μM, 24 h). Membranes were probed for GFP or α-tubulin. Quantification of CTF:ICD and α-tubulin band intensities is shown underneath. Data are mean ± SD (n = 4 – 5, N = 3 extractions; unpaired t-test). (C) Representative Na^+^ currents in MDA-MB-231-β1-GFP cells pre-treated with DMSO (0.01 % (v/v), 24 h) or DAPT (1 μM, 24 h), following depolarisation between -80 mV and +30 mV for 250 ms from -120 mV. Peak current density (PCD) is compared on the right (unpaired t-test). Data are mean ± SD (n= 8, N = 3). (D) Western blot of lysates from MDA-MB-231-β1-GFP cells treated with DMSO (0.1% (v/v), 24 h), L685,458 (10 μM, 24 h) or Avagacestat (10 μM, 24 h). Membranes were probed for GFP or α-tubulin. Quantification of CTF/ICD band intensity is shown underneath. Data are mean ± SD (N = 3 extractions; one-way ANOVA with Dunnett’s test). (E) Representative Na^+^ currents in MDA-MB-231-β1-GFP cells pre-treated with DMSO (0.1 % (v/v), 24 h) or Avagacestat (10 μM, 24 h). PCD is shown underneath. Data are mean ± SD (n = 7 - 8, N = 3, unpaired t-test). (F) PCD in MDA-MB-231-GFP cells and MDA-MB-231-β1-GFP cells following pre-treatment with DMSO (n = 18, N = 3), 1 μM L-685,458 (n = 12, N = 3) and 10 μM L-685,458 (n = 7, N = 3), one-way ANOVA with Tukey’s test. (G) Time taken for half-maximal recovery from inactivation for MDA-MB-231-β1-GFP cells pre-treated with DMSO (0.01 – 0.1 % (v/v), 24 h), DAPT (1 μM, 24 h), Avagacestat (Ava, 10 μM, 24 h) or L,685-458 (1 μM, 24 h; n = 8, N = 3). Cells were depolarised to 0 mV for 25 ms, then held at -120 mV for *t* s before second depolarisation to 0 mV. *t* ranged from 1-500 ms. Data are mean ± SD (unpaired t-test). ns = not significant, * = P < 0.05, ** = P < 0.01.

### Effect of γ-secretase inhibition on β1-GFP subcellular distribution

We next attempted to establish the subcellular site of secretase cleavage of β1-GFP. γ-secretase is present throughout the cell, including the plasma membrane (55), nuclear envelope (56), endoplasmic reticulum (57), mitochondria (58), Golgi apparatus (59), endosomes (60), and lysosomes (61). Interestingly, following DAPT treatment, the Pearson’s correlation coefficient (PCC) between β1-GFP and EEA1 decreased by ∼10 % (P < 0.05; n = 25–26; Mann-Whitney U-test; Figure 5A, C), suggesting a decrease in endosomal β1-GFP following γ-secretase inhibition. There was no difference in the PCC between β1-GFP and LAMP1 in vehicle- vs. DAPT-treated cells (P = 0.44; n = 23 – 27; Mann-Whitney U-test; Figure 5B, C), suggesting lysosomes may not be a site of secretase processing for β1-GFP. The observations that DAPT reduced the endosomal (but not lysosomal) β1-GFP level, and that the majority of β1-GFP is retained intracellularly (Figure 2), suggest that some β1-GFP may be trafficked to lysosomes for degradation without being expressed at the surface, as has been shown for amyloid precursor protein (APP) (62).

**Figure 5.**
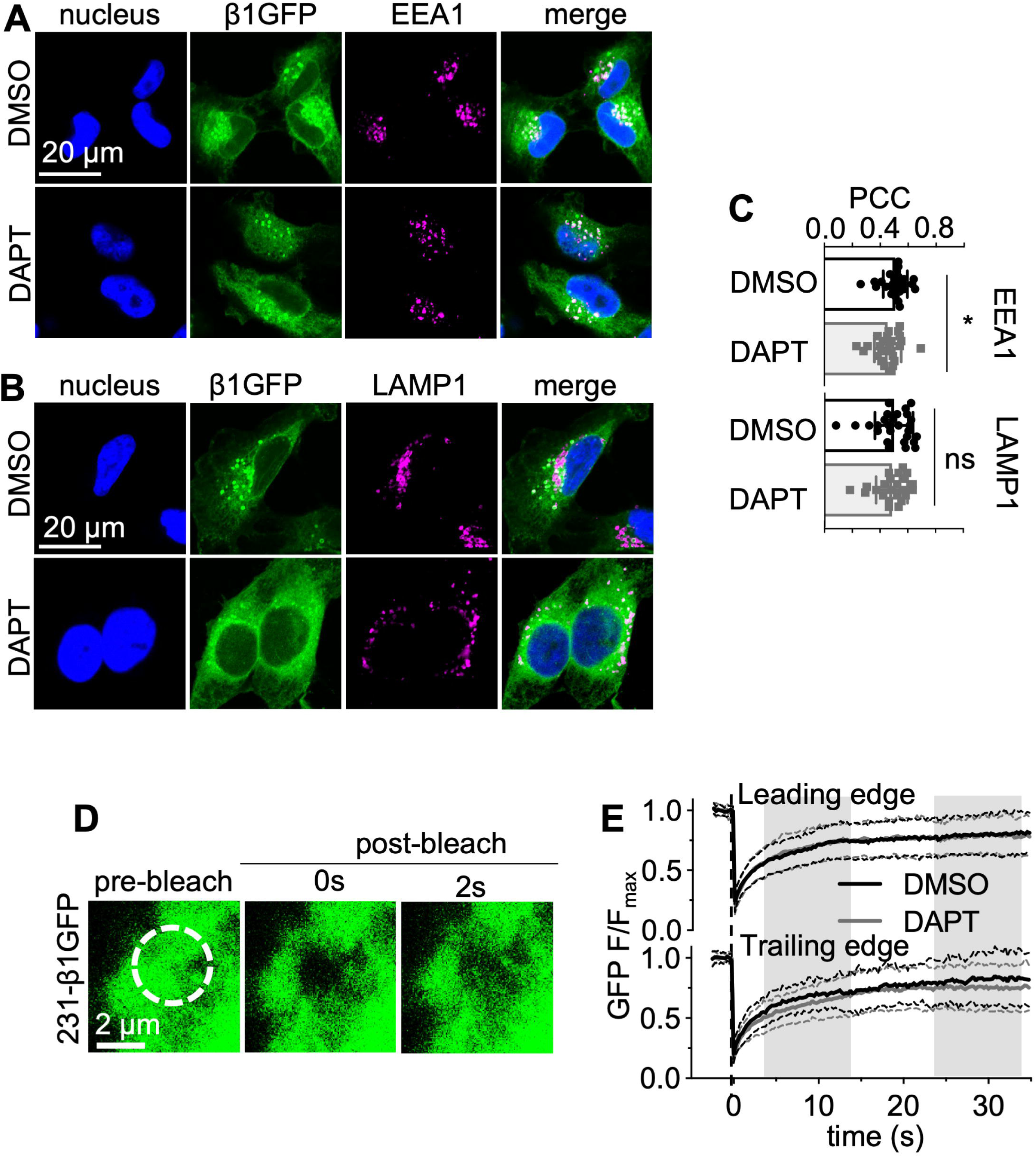
Effect of γ-secretase inhibition on β1-GFP localisation. MDA-MB-231-β1GFP cells were treated with DMSO (0.01 % (v/v), 24 h) or DAPT (1 μM, 24 h) and stained for EEA1 (A, magenta) or LAMP1 (B, magenta) and DNA (DAPI, blue). Endogenous GFP is shown in green. (C) Pearson’s correlation coefficient (PCC) for co-localisation of GFP and EEA1 or LAMP1. Data are mean ± SD (n = 23 - 27, N = 3), Mann Whitney U-test, ns = not significant, * = P<0.05, ** = P<0.01. (D) Representative recovery of GFP fluorescence at the leading edge of a live MDA-MB-231-β1-GFP cell following photobleaching with a 488 nm laser (40iterations, 100 % laser power). (E) Quantification of the recovery of GFP fluorescence in MDA-MB-231-β1-GFP cells at the leading and trailing edges following treatment with DMSO (0.01 % (v/v), 24 h, black line) or DAPT (1 μM, 24 h, grey line). Data are mean (solid line) ± SD (dotted line), n = 25 - 28, N = 3.

To obtain a better spatiotemporal understanding of γ-secretase-mediated cleavage of β1-GFP, we next followed GFP distribution in live MDA-MB-231-β1-GFP cells using confocal microscopy and fluorescence recovery after photobleaching (FRAP). Regions of interest (ROIs) were photobleached at the leading and trailing edges of control and DAPT-treated MDA-MB-231-β1-GFP cells (Figure 5D, E). Interestingly, no differences in the proportion of GFP that was freely mobile, or the time taken for half-maximal fluorescence recovery, were detected at the leading or trailing edges (Table 1). This result suggests that DAPT treatment had no effect on spatiotemporal cycling dynamics of β1-GFP.

**Table 1.**
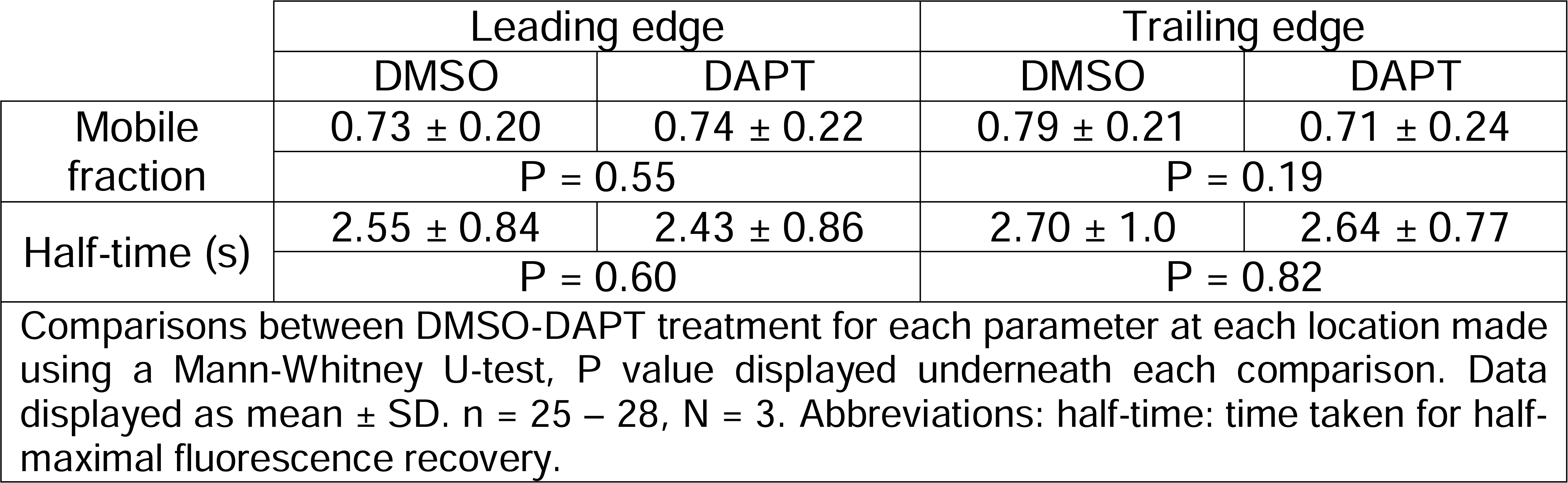
Mobility parameters of β1-GFP at the leading and trailing edges of MDA-MB-231 cells following DAPT treatment

### Functional consequences of β1ICD-GFP overexpression

To further study the functional activity of the β1-ICD, we over-expressed β1ICD-GFP in MDA-MB-231 cells and measured its effect on Na^+^ current using patch clamp recording (Figure 6A). Interestingly, β1-GFP-expressing cells (-11.72 ± 3.56 pA/pF) and β1ICD-GFP-expressing cells (-9.33 ± 4.15 pA/pF) displayed significantly larger Na^+^ current density compared to control GFP-expressing cells (-4.38 ± 2.49 pA/pF; P < 0.01; n = 14 - 16; one-way ANOVA; Figure 6B). In addition, β1ICD-GFP accelerated the recovery from inactivation to the same extent as β1-GFP (P < 0.05; n = 10; one-way ANOVA; Figure 6C).

**Figure 6.**
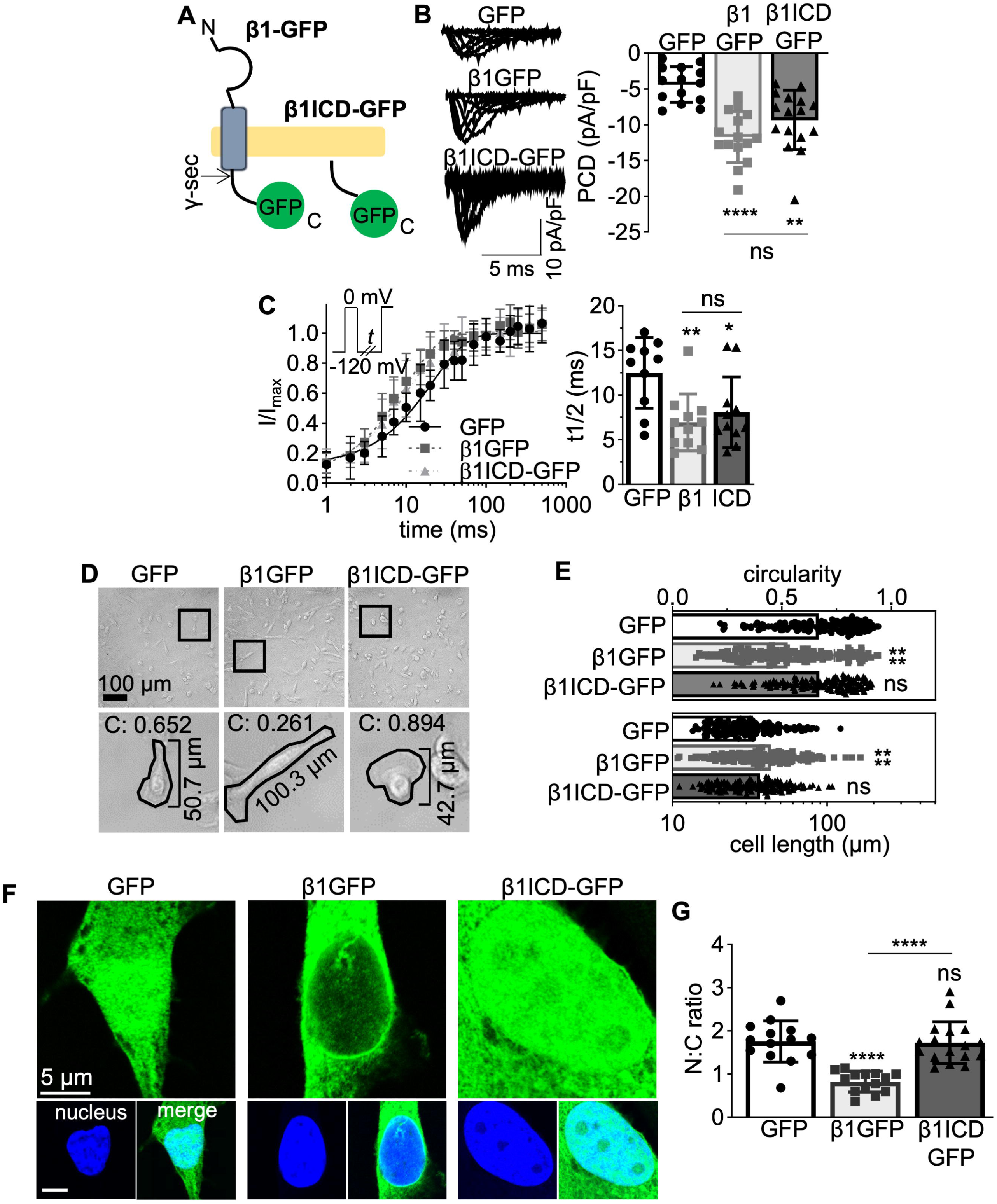
Function and subcellular localisation of β1ICD-GFP. (A) Schematic of full-length β1-GFP alongside β1ICD-GFP. (B) Representative whole-cell Na^+^ currents in MDA-MB-231-GFP, MDA-MB-231-β1-GFP and MDA-MB-231-β1ICD-GFP cells, following depolarisation between -80 mV and +30 mV, for 250 ms, from -120 mV. Peak current density (PCD) is shown on right. Data are mean ± SD (n =14 – 16, N = 3), Kruskal-Wallis test. (C) Recovery from inactivation and time taken for half-maximal recovery from inactivation (t1/2) Cells were depolarised to 0 mV for 25 ms, then held at -120 mV for *t* s before re-stimulation at 0 mV. *t* ranged from 1-500 ms. Data are mean ± SD (n = 8, N = 3). (D) Brightfield images of MDA-MB-231-GFP, MDA-MB-231-β1-GFP and MDA-MB-231-β1ICD-GFP cells at 20x magnification. Bottom row displays zoomed images from boxes above, with masks over example cells depicting circularity index (“C”) and cell length. (E) Quantification of circularity index and cell length from brightfield images like those in D. Data are mean ± SD (n = 150, N = 3), Kruskal-Wallis test. (F) Nuclear images of the GFP signal within fixed MDA-MB-231 cells expressing GFP, β1-GFP or β1ICD-GFP. Main panels show GFP signal (green), sub-panels depict DAPI (blue) and DAPI/GFP merge. (G) Nuclear:cytoplasmic signal density ratio (N:C ratio). Data are mean ± SD (n = 14 – 17, N = 3), one-way ANOVA with Tukey’s test. ns = not significant, * = P < 0.05, ** = P < 0.01, **** = P < 0.0001.

β1-GFP over-expression in MDA-MB-231 cells increases process length and decreases process width (39). We therefore next tested the effect of β1ICD-GFP on cellular morphology. β1ICD-GFP overexpression had no effect on cell length (P = 0.14; n = 150; Kruskal-Wallis test; Figure 6D, E) or circularity (P = 0.99; n = 150; Kruskal-Wallis test; Figure 6D, E), relative to expression of GFP alone. In contrast β1-GFP increased cell length by ∼30 % (P < 0.0001; n = 150; Kruskal-Wallis test; Figure 6D, E) and reduced circularity by ∼27 % relative to GFP control (P < 0.0001; n = 150; Kruskal-Wallis test; Figure 6D, E). Together, these data suggest that β1ICD-GFP recapitulates the electrophysiological effects of full-length β1-GFP on Na^+^ current in MDA-MB-231 cells but does not itself promote changes in cellular morphology.

### Subcellular distribution of β1ICD-GFP

To determine the extent of β1-ICD localisation to the nucleus, we imaged β1ICD-GFP-, β1-GFP- and GFP-expressing cells by confocal microscopy (Figure 6F). We used GFP-expressing cells as a control for stochastic movement of small proteins because GFP is known to diffuse throughout the cell, including into the nucleus (63). The nuclear signal for β1ICD-GFP was higher than for β1-GFP, consistent with not all full-length β1-GFP being cleaved at steady state (P < 0.0001; n = 14–17; one-way ANOVA; Figure 6G). In addition, GFP and β1ICD-GFP had similar nuclear:cytoplasmic signal density ratio (P = 0.98; Figure 6G). These data suggest that β1ICD-GFP is present within the nucleus. However, it is possible that nuclear localisation of β1ICD-GFP may be due to stochastic diffusion from the cytoplasm, similar to the case with GFP.

To more accurately evaluate whether β1ICD-GFP distribution differs from GFP, we compared the mobility of both proteins using FRAP. Initially, a ROI within the cytoplasm of MDA-MB-231-GFP and MDA-MB-231-β1ICD-GFP cells was photobleached and fluorescence recovery measured (Figure 7A, B). GFP and β1ICD-GFP displayed similar mobility kinetics within the cytoplasm, with both proteins having a comparable mobile fraction of ∼1 (P = 0.07; n = 15; Mann-Whitney U-test; Figure 7C) and time taken for half maximal fluorescence recovery (P = 0.13; n = 15; Welch’s t-test; Figure 7D). These data suggest that β1ICD-GFP and GFP have similar spatiotemporal expression within the cytoplasm of MDA-MB-231 cells.

**Figure 7.**
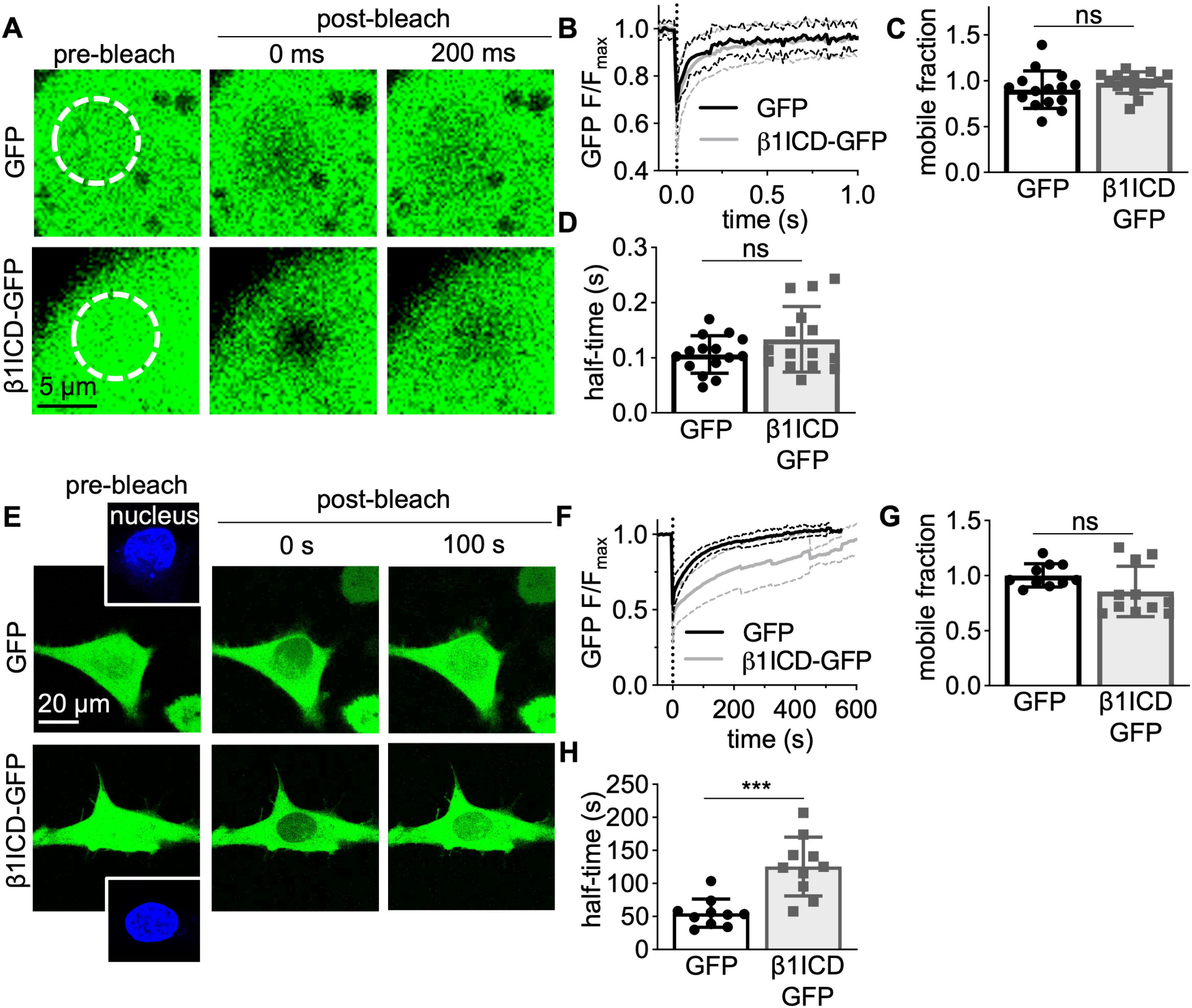
Intracellular mobility of β1ICD-GFP. (A) Live-cell confocal imaging of MDA-MB-231-GFP (top row) and MDA-MB-231-β1ICD-GFP (bottom row) cells in the cytoplasm. Cells were imaged every 14 ms for 2 s and a 1.5 - 2 μm wide region-of-interest photobleached with a 488 nm laser (100 % laser power, 40 iterations). Images displayed are immediately prior to photobleaching (first column), immediately following photobleaching (second column) and 200 ms after photobleaching (third column). (B) Fluorescence recovery within the region of interest in the cytoplasm (n = 15, N = 3). Data are mean (solid line) ± SD (dotted line). (C) Quantification of the mobile fraction in the cytoplasm (n = 15, N = 3). (D) Time taken for half-maximal fluorescence recovery (half-time, s) in the cytoplasm (n = 15, N = 3). (E) Live-cell confocal imaging of MDA-MB-231-GFP (top row) and MDA-MB-231-β1ICD-GFP (bottom row) cells in the nucleus. Cells were imaged every 250 ms and photobleached with a 488 nm laser (100 % laser power, 40 iterations). Time series were acquired until five successive images without an increase in nuclear fluorescence were acquired. Images displayed are immediately prior to photobleaching (first column), immediately following photobleaching (second column) and 100 s after photobleaching (third column). Nuclei were photobleached following masking using the Hoechst 33342 signal (shown in sub-panels in the first column, blue). (F) Fluorescence recovery within the region of interest in the nucleus (n = 10, N = 3). Data are mean (solid line) ± SD (dotted line). (G) Quantification of the mobile fraction in the nucleus (n = 10, N= 3). (H) Time taken for half-maximal fluorescence recovery (half-time, s) in the nucleus (n = 10, N = 3). Data are mean ± SD. Unpaired t-tests were used to test significance, except Mann-Whitney U-test for H. ns = not significant, *** = P < 0.001.

Next, we compared the nuclear import kinetics of GFP and β1ICD-GFP. Other cleaved ICDs, such as Notch, are trafficked to the nucleus as part of a heteromeric complex (64). Such a complex is expected to have slower import kinetics than soluble GFP, which can diffuse directly through nuclear pores. We labelled live cells with the nuclear dye, Hoechst 33342, and then photobleached the overlapping nuclear GFP fluorescence and measured fluorescence recovery over time (Figure 7E, F). Both GFP and β1ICD-GFP demonstrated a similar mobile fraction of ∼ 1 (P = 0.08; n = 10; unpaired t-test; Figure 7G), suggesting negligible immobilised protein is present within the nucleus. However, the time taken for half-maximal fluorescence recovery was ∼2.5 fold greater for β1ICD-GFP relative to GFP (P < 0.001; n = 10; Mann-Whitney U-test; Figure 7H), implying different nuclear import kinetics for both proteins. According to the experimentally verified cubic relationship between molecular weight and time taken for nuclear import, the five additional kilodaltons of β1ICD-GFP should increase diffusion time through nuclear pores by ∼70 % compared to GFP (65). Therefore, β1ICD-GFP may be entering the nucleus as part of a larger protein complex.

### Effect of β1-GFP and β1ICD-GFP expression on tetrodotoxin sensitivity

To further evaluate the involvement of the ICD in regulating Na^+^ current, we over-expressed β1STOP-GFP, which lacks the ICD (66), in MDA-MB-231 cells (Figure 8A). β1STOP-GFP did not significantly increase peak current density compared to GFP (P = 0.89; n = 15; one-way ANOVA; Figure 8B). This result underscores the importance of the β1-ICD for increasing Na^+^ current density. *SCN5A* mRNA constitutes 82 % of total α-subunit-encoding mRNA present and the Na^+^ current in MDA-MB-231 cells is predominantly TTX-resistant, suggesting Na_v_1.5 is the main α-subunit at the cell surface (50). Given that β1-GFP and β1ICD-GFP increased Na^+^ current, we used TTX to examine whether the composition of functional α subunits at the plasma membrane was altered, as 1 µM TTX blocks Na_v_1.1–1.4, Na_v_1.6, and Na_v_1.7, but not Na_v_1.5. When perfused with 1 μM TTX, the Na^+^ current in MDA-MB-231-GFP cells was not significantly altered (P = 0.59; n = 9; one-way ANOVA; Figure 8C, left panel), consistent with a TTX-resistant channel, most likely Na_v_1.5, predominant at the cell surface. In MDA-MB-231-β1-GFP cells, however, 1 µM TTX significantly decreased the Na^+^ current by 33 ± 2 % (P < 0.0001; n = 9; one-way ANOVA; Figure 8C, centre panel). In addition, 1 µM TTX application also decreased Na^+^ current in β1ICD-GFP-expressing cells by 35 ± 3 % (P < 0.0001; n = 9; one-way ANOVA; Figure 8C, right panel). These data suggest that β1-GFP and β1ICD-GFP are both capable of increasing the proportion of TTX-sensitive α subunits at the plasma membrane.

**Figure 8.**
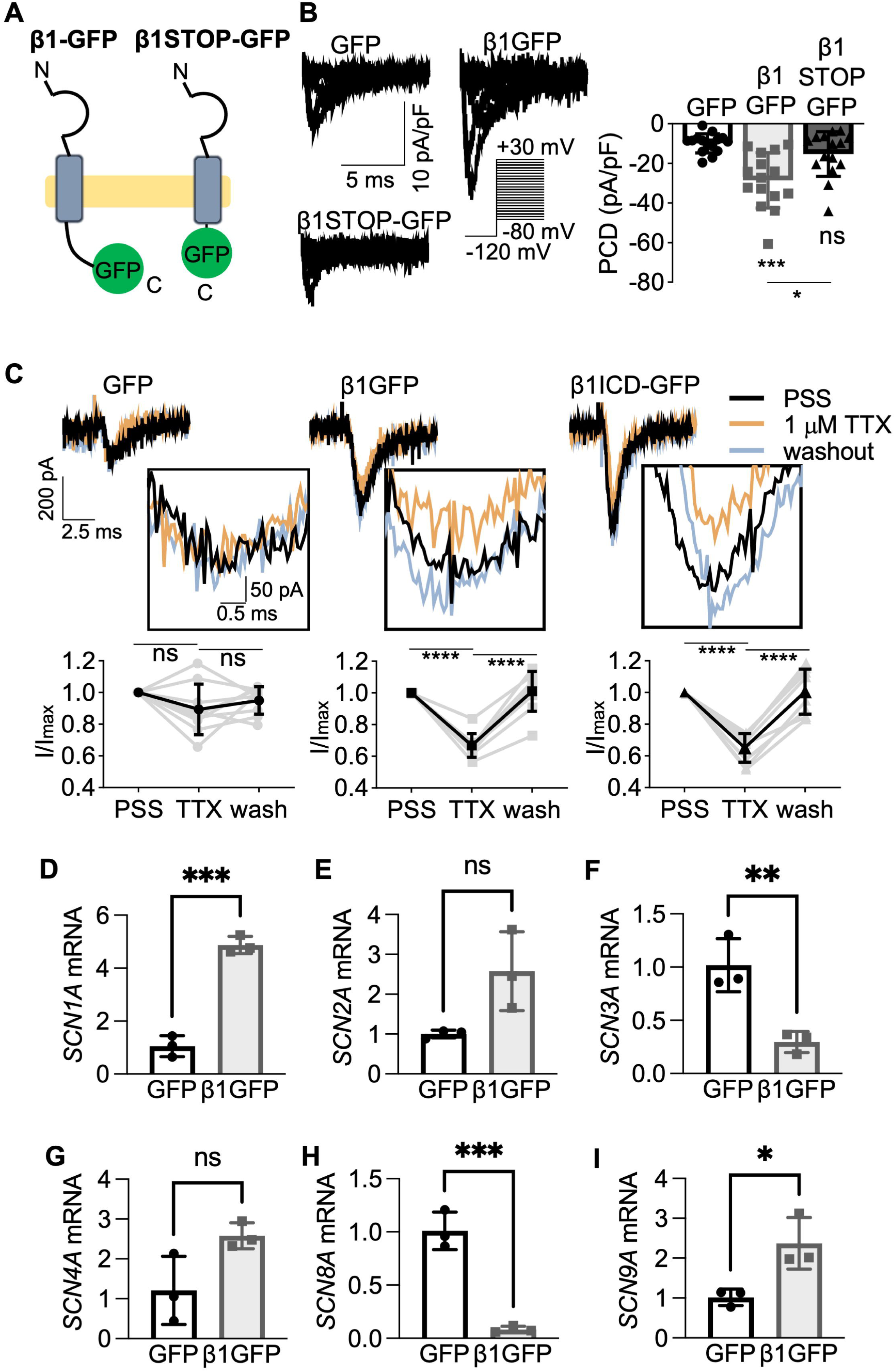
Regulation of TTX-sensitive Na^+^ current by β1-GFP and β1ICD-GFP. (A) Schematic of full-length β1-GFP alongside β1STOP-GFP. (B) Representative whole-cell Na^+^ currents generated following depolarisation between -80 mV and +30 mV, for 250 ms, from - 120 mV. Peak current density (PCD) is shown on right. Data are mean ± SD (n = 15, N = 3), Kruskal-Wallis test. (C) Representative traces of I_Na_ in MDA-MB-231 cells expressing GFP, β1-GFP, or β1ICD-GFP in standard recording solution (PSS, black line), 1 μM TTX (orange line) and following washout (grey line). Quantification of the reduction in peak current density in 1 μM TTX and following washout is shown underneath, normalised to PSS. (D-I) Relative mRNA levels (fold change compared to GFP control) for *SCN1A* (D), *SCN2A* (E), *SCN3A* (F), *SCN4A* (G), *SCN8A* (H), and *SCN9A* (I). Data are mean ± SD (n = 6 - 9, N = 3 for A-C; N = 3 for D-I), repeated measures ANOVA for A-C, t-test for D-I. ns = not significant, * = P < 0.05, ** = P < 0.01, *** = P < 0.001, **** = P < 0.0001.

Given that β1-ICD has been shown to translocate to the nucleus and regulate gene transcription (46), we next evaluated the effect of β1-GFP on TTX-sensitive α-subunit mRNA levels in MDA-MB-231 cells, compared to GFP. Overexpression of β1-GFP significantly increased the mRNA level of *SCN1A* (P < 0.001; n = 3; t-test; Figure 8D) and *SCN9A* (P < 0.05; n = 3; t test; Figure 8I). There was also a small increase in *SCN2A* and *SCN4A* expression, although this was not statistically significant (P = 0.052 and P = 0.061, respectively; n = 3 for both; Figure 8E, G). Finally, there was a significant reduction in *SCN8A* expression (P < 0.001; n = 3; t-test; Figure 8H). However, given that *SCN8A* has previously been shown to be expressed in a truncated form in MDA-MB-231 cells (50), this reduction is unlikely to be physiologically relevant. We therefore conclude that the elevated TTX-sensitive Na^+^ current present in β1-GFP cells is likely carried by Na_v_1.1 and/or Na_v_1.7 and that the regulation of these subunits by β1-GFP may, at least in part, be transcriptional.

## Discussion

In this study, we show that both β1-GFP and its γ-secretase cleavage product, β1ICD-GFP, are functionally active in MDA-MB-231 breast cancer cells. We show that the majority of β1-GFP is retained intracellularly, specifically within the endoplasmic reticulum, Golgi apparatus, endosomes, and lysosomes. A reduction in endosomal β1-GFP level occurred following γ-secretase inhibition, implicating endosomes, and/or the preceding plasma membrane, as an important site for secretase processing. A small fraction of β1-GFP was detected in the nucleus, and soluble β1ICD-GFP demonstrated unique nuclear expression and import kinetics. Furthermore, β1ICD-GFP was necessary and sufficient to increase Na^+^ current measured at the plasma membrane. Finally, both β1-GFP and β1ICD-GFP increased TTX sensitivity of the Na^+^ current. We therefore propose that the proteolytically released β1-ICD is a critical regulator of α subunit function and expression in cancer cells. A strength of our study compared with previous studies (45,46,48), is that, by using GFP-tagged constructs, we were able to visualise β1/β1-ICD dynamics in live cells. However, a caveat with this approach is that we cannot exclude the possibility that the GFP tag may interfere with function of the native protein and its cleavage products. Nonetheless, our findings are generally consistent with these other reports, suggesting that any disruption of β1 function by the GFP tag may be minor.

The requirement for β1 insertion into the plasma membrane as a prerequisite for secretase processing is dependent on S-palmitoylation of a cysteine residue found at the transmembrane-intracellular interface of β1 (47). Our work supports this finding and suggests that although secretase cleavage is not necessary for the β1-dependent increase of Na^+^ current *per se*, presence of the β1-ICD is required. Previous work has demonstrated that the extracellular domain of β1 is sufficient to accelerate channel inactivation in *Xenopus* oocytes (40,67), and more recent voltage clamp fluorimetry and protein crystallography studies have further shown interaction between β1 and voltage-sensing domains of α subunits (8,42,43). Thus, a mechanistic understanding of how β1 modulates α subunit gating and kinetics is becoming clearer. However, the mechanism underlying β1-mediated potentiation of VGSC expression at the plasma membrane and the consequent increase in current density remains uncertain. An interaction site within the β1-ICD is required to increase surface expression of Na_v_1.1 and Na_v_1.2 in heterologous cell models (4,41). Furthermore, the β1-ICD has been shown to increase Na_v_1.8 current density through deletion and β1-β2 chimera experiments (68). Soluble, intracellular non-pore-forming subunits for other ion channel families also exist. For example, various Ca_v_β, K_v_β and K^+^ channel interacting protein (KChIP) subunits can modulate channel activation/inactivation kinetics and increase surface expression (in the case of Ca_v_βs and KChIPs) (69-72). Furthermore, the APP-ICD has been shown to increase Na_v_1.6 current when co-expressed in *Xenopus* oocytes (73). Together, our data support the notion that the β1-ICD is a functionally active regulator of α subunit expression at the plasma membrane of MDA-MB-231 cells. However, the precise involvement of γ-secretase cleavage on these mechanisms appears complex, given that full-length β1-GFP and β1ICD-GFP both promote Na^+^ current, and that pharmacological inhibition of γ-secretase activity had no effect. Furthermore, the β1-mediated targeting of α subunits to the plasma membrane may be cell type-specific and the over-expression systems used may not accurately reflect the stoichiometric balance that occurs between endogenous α and β subunits and/or their subcellular localization. We previously showed that endogenous β1 is present in subcellular compartments in breast cancer cells (74). Further work is required to elucidate the context-dependent localization and trafficking of endogenous β1 subunits in different cell types.

Proteolytic processing is also an important regulator of the CAM function of β subunits in neurons and cancer cells. Pharmacological blockade of γ-secretase cleavage inhibits β1-mediated neurite outgrowth in cerebellar granule neurons (3). Additionally, γ-secretase inhibition decreases β2-induced transcellular adhesion and cell migration (45). The soluble cleaved extracellular Ig domains of β1 and β4, as well as the soluble splice variant β1B, have been shown to promote neurite outgrowth (24,25,75). In MDA-MB-231 cells, β1-GFP, but not a mutant lacking the Ig domain, promotes neurite-like process outgrowth (38). In agreement with these observations, we found here that, in contrast to full-length β1-GFP, β1ICD-GFP was not capable of promoting process outgrowth in MDA-MB-231 cells. Thus, regulated proteolysis at the plasma membrane is likely to be a key mechanism by which the CAM function of β1 is modulated to fine-tune neurite outgrowth, neuronal pathfinding, fasciculation and cell migration (15,19-24,47). Interestingly, endogenous β1 expression is higher in MCF-7 cells than in MDA-MB-231 cells (39), although γ-secretase activity has been reported in both breast cancer cell lines (76). β1 expression in breast cancer cells has been shown to increase adhesion *in vitro* and promote neurite-like process outgrowth, tumour growth, and metastasis *in vivo* (38,39). These effects generally fit with emerging data indicating that expression/activity of VGSCs promotes invasion and metastasis across multiple cancer types where these channels have been shown to be expressed (77). Further work is required to establish whether variation in endogenous β1 expression between different cancer cell types may determine the impact of γ-secretase activity on β1 function.

A number of secretase-cleaved ICDs have been shown to translocate to the nucleus and are involved in gene regulation, such as the ICDs for APP (78), CD44 (79) and Notch (80). The ICDs of β1 and β2 have been shown to accumulate in the nucleus of heterologous cells (46,48). In addition, the β2-ICD has been shown to upregulate *SCN1A*/Na_v_1.1 mRNA and protein expression (48), raising the possibility that secretase cleavage may modulate α subunit expression via altering α subunit transcription. Moreover, the β1-ICD has recently been shown to regulate the transcription of multiple genes (46). Our findings provide additional insight with respect to putative β1-ICD nuclear function, showing accumulation in the nucleus of live and fixed cells. In addition, although the endogenous Na^+^ current in these cells is carried by Na_v_1.5 (50,51), we observed an increase in *SCN1A* and *SCN9A* mRNA expression and TTX-sensitive Na^+^ current, suggesting increased expression and/or trafficking of these α subunits in the presence of β1-GFP. Further work is needed to determine how the β1-ICD is directed to the nucleus and the mechanism by which it regulates gene expression.

Taken together with the available literature (4,38,39,41,46-48), our data suggest that the β1-ICD promotes plasma membrane α subunit expression in breast cancer cells via several mechanisms (Figure 9). The observation that β1STOP-GFP failed to increase Na^+^ current further identifies a requirement for the β1-ICD. β1STOP-GFP did, however, accelerate recovery from inactivation, suggesting that the β1-ICD is not required for regulation of α subunit inactivation kinetics. Therefore, it appears that the mechanisms underlying increased Na^+^ current density, presumably via increased plasma membrane expression, and channel inactivation are distinct. Our data suggest that processing by γ-secretase may play a role in regulating β1 function in breast cancer cells, adding to emerging evidence in other cell systems (46). The important role of γ-secretase activity in cancer progression (49), together with the growing evidence suggesting that β1ICD-mediated cellular changes promote pathologies including epilepsy, cardiac arrhythmia and cancer (46), highlight the significance of this signalling axis to pathophysiologies associated with abnormal β1 function. Further work is required to address the generalizability of these observations across different disease states.

**Figure 9.**
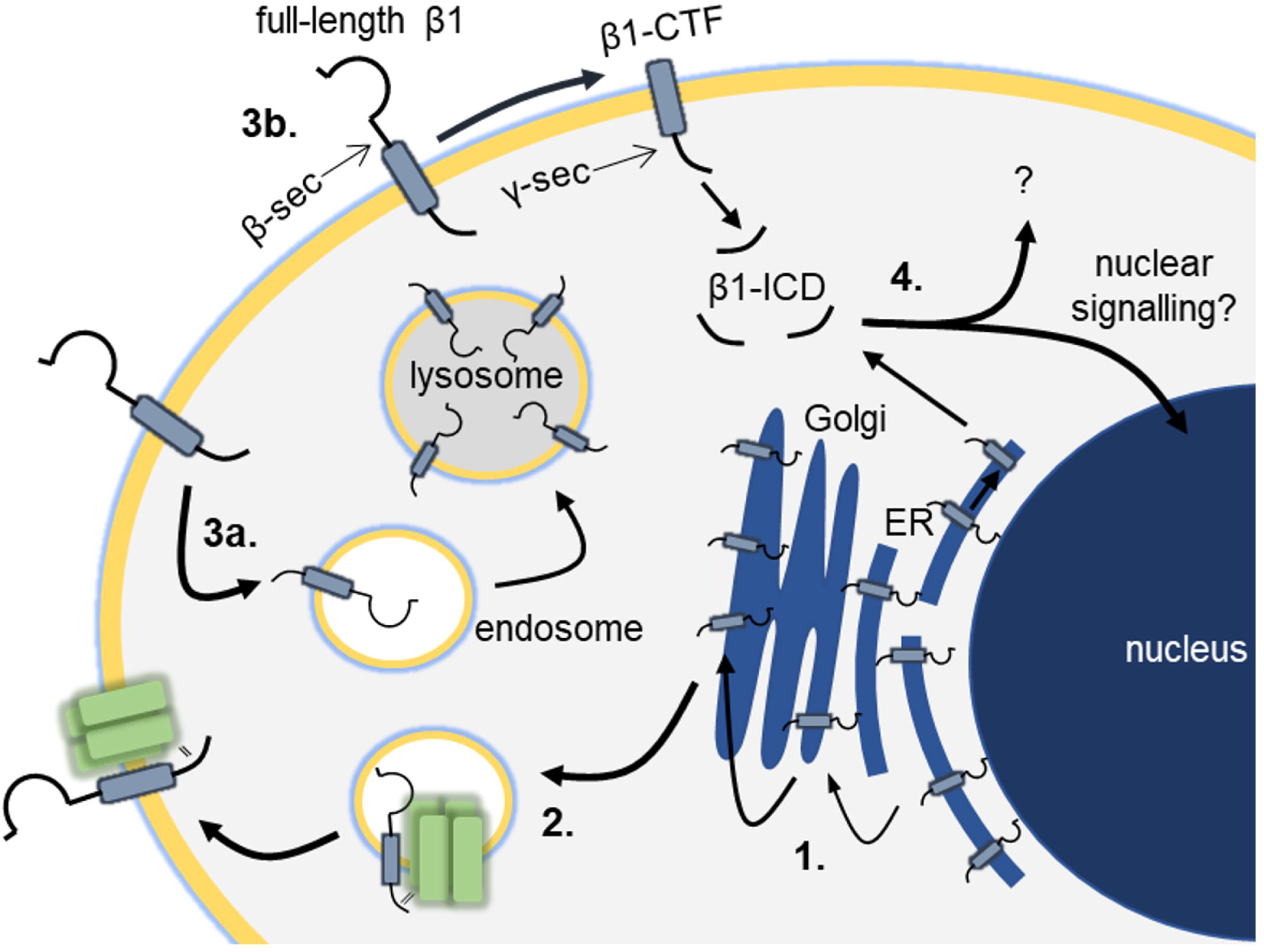
Model for β1 cycling in MDA-MB-231 cells. 1. β1 progresses through the endoplasmic reticulum and Golgi apparatus. The majority of cellular β1 is sequestered within the ER. 2. A fraction of β1 is trafficked to the cell surface associated by its ICD to an α-subunit. 3. β1 is either internalised (3a) or processed by secretases at the plasma membrane (3b). 4. Cleaved β1-ICD may be degraded or translocate to the nucleus to regulate gene expression.

## Experimental Procedures

### Cell culture

Human mammary carcinoma MDA-MB-231 cells were maintained in Dulbecco’s Modified Eagle Medium (DMEM) supplemented with 5 % (v/v) foetal bovine serum and 4 mM L-glutamine and cultured at 37 °C/ 5 % CO_2_ (74). Stable MDA-MB-231-GFP and MDA-MB-231-β1-GFP cell lines were generated previously (39). Cell culture medium was supplemented with G418 (200 μg/ml, Sigma) for MDA-MB-231-GFP cells or Hygromycin B (100 μg/ml, Invitrogen) for the other transfected cell lines.

### Pharmacology

γ-secretase inhibitors used in this study were as follows: avagacestat (10 μM, 24 h treatment time, Sigma), N-[(3,5-Difluorophenyl)acetyl]-L-alanyl-2-phenyl]glycine-1,1-dimethylethyl ester (DAPT) (1 μM, 24 h, Santa Cruz Biotech) and L-685,458 (1 - 10 μM, 24 h, Santa Cruz Biotech). DMSO was the vehicle for all drugs and DMSO concentration did not exceed 1:1000 at working concentration. Chloroquine (Tokyo Chemicals Industry) was dissolved in H_2_O and used at 10 μM for 24 h.

### Site-directed mutagenesis

A pEGFPN-1 expression plasmid encoding *Rattus norvegicus* β1, with a C-terminal enhanced GFP tag, was developed previously (39). The insert encoding β1-GFP was sub-cloned into a pcDNA3.1 expression vector (Invitrogen), following digest of 1 μg of both plasmids with 1 U of FastDigest NheI (Thermo) and FastDigest NotI (Thermo) for 30 min at 37°C. Fragments were gel purified (following kit instructions, Macherey-Nagel) and 30 ng of insert and 10 ng of vector ligated using 6U of T4 DNA ligase (Thermo, 1h, room temperature). pcDNA3.1-β1-GFP was used to produce several mutant β1 constructs using PCR-based site-directed mutagenesis following manufacturer’s instructions (Phusion site-directed mutagenesis kit, Thermo) (38). β1 constructs produced were as follows: β1ICD-GFP (sequence starting at Tyr163 [mature protein amino acid numbering used, i.e. after 19 amino acid signal peptide cleaved]) and β1STOP-GFP (β1-ICD deletion, sequence terminated after Lys165), used previously (66). Primers used were as follows: β1ICD-GFP forward-AAGAAGATTGCTGCTGCCACG and β1ICD-GFP reverse-CATCTTGGGTCTCCCTATAGTGAGTCGTATTA (annealing temperature, T_A_ – 69 °C). β1STOP-GFP forward – CGAATTCTGCAGTCG and β1STOP-GFP reverse – CTTCTTGTAGCAGTACAC (T_A_ – 60 °C). All construct sequences were confirmed by Sanger sequencing (Source Bioscience).

### Transfection and generation of stably transfected cell lines

MDA-MB-231 cells were transfected using jetPRIME (Polyplus) at a DNA:jetPRIME ratio (ng:nl) of 1:2. Medium was changed after 4 h. Stable cell lines were produced following Hygromycin B treatment (300 μg/ml until non-transfected cells died) then ring selection and propagation of single colonies, maintained in Hygromycin B (100 μg/ml).

### Protein extraction and western blot

Protein extraction and western blotting were carried out as described previously, with some modifications (38). Total protein was extracted from a confluent 15 cm dish of cells and suspended in 50 mM Tris, 10 mM EGTA, with protease inhibitors (Roche). Lysates were diluted in Laemmli buffer at a ratio of 4:1 and heated at 80 °C for 10 min. Protein (30 – 100 μg) was separated by sodium dodecyl sulphate-polyacrylamide gel electrophoresis (12 % acrylamide, 120 V, 2 h) and transferred onto a nitrocellulose (1.3 A, 25 V, 10 min) or a PVDF (2.5 A, 25 V, 3 min) membrane by semi-dry transfer. Primary antibodies were rabbit anti-GFP (ab6556, 1:2500, Abcam) or mouse anti-α-tubulin (clone DM1A, 1:10,000, Sigma). Secondary antibodies were HRP-conjugated goat anti-mouse (1:1000, Thermo Scientific) or goat anti-rabbit (1:1000, Thermo Scientific). Chemiluminescence was detected using an iBRIGHT imaging system (Invitrogen) or X-ray film (Fujifilm) following West Dura application (5 min, Thermo Scientific). Densitometry was performed on western blot bands to estimate protein quantity using ImageJ 1.51i (81).

### Morphology assay

10,000 cells were seeded into a well of a 24-well plate and left for 72 h prior to image acquisition. Cells were fixed using 4 % (w/v) paraformaldehyde (PFA) in PBS at room temperature for 10 min and washed with 0.1 M phosphate buffer (PB, 81 mM Na_2_HPO_4_, 19 mM NaH_2_PO_4_, pH 7.4). Five brightfield images of each well were acquired, as well as GFP images to ensure construct expression. Images were exported to ImageJ for analysis. Cell morphology was assessed by manually masking the first 50 randomly selected cells and measuring circularity and Feret’s diameter (estimator of cell length) using the in-built analysis ImageJ plugin. The experiment was repeated three times.

### Immunocytochemistry

Protocols were adapted from (15,82). Cells grown on 13 mm glass coverslips were fixed with 4 % (w/v) PFA (dissolved in PBS) for 5 min and washed three times in 0.1 M PB. In some experiments, cells were incubated in 50 μg/ml digitonin (Santa Cruz Biotechnology) for 15 min and Triton X-100 was omitted in the subsequent steps (53). Depending on the primary antibody, cells were blocked for 1 h in either PBTGS (0.3 % (v/v) Triton X-100 and 10 % (v/v) normal goat serum in 0.1 M PB) for anti-Lamin B2 and anti-GFP antibodies, or BPS (0.5 % (w/v) BSA and 0.05 % (w/v) saponin dissolved in 0.1 M PB) for all other antibodies. Primary antibody (diluted in blocking solution) was applied for 1 h (antibodies diluted in BPS) or overnight (antibodies diluted in PBTGS). The following primary antibodies were used: anti-GFP (mouse, Neuromab, clone N86/38, 1:1000), anti-Lamin B2 (mouse, Invitrogen, clone E-3, 1:500), anti-EEA1 (mouse, clone 14/EEA1, BD Bioscience, 1:500), anti-LAMP1 (mouse, Biolegend, clone H4A3, 1:1000), anti-calnexin (mouse, BD Bioscience, clone 37/CNX, 1:50) and anti-TGN46 (rabbit, Proteintech, 1:1000). Cells were then incubated in goat anti-mouse Alexa Fluor 568, or goat anti-rabbit Alexa Fluor 568 or 647 (1:500 in blocking solution, Thermo) for 1 h (antibodies diluted in BPS) or 2 h (antibodies diluted in PBTGS). Cells were incubated in 500 ng/ml 4’,6-diamidino-2-phenylindole (DAPI, diluted in 0.1 M PB, Sigma, 10 min) prior to mounting using Prolong Gold (Invitrogen).

### Confocal microscopy

Images were acquired using a Zeiss LSM 880 laser-scanning confocal microscope with Airyscan technology, using a Plan-Apochromat 63x oil immersion objective lens (NA = 1.4), controlled by ZEN2 software. The pinhole was set to 1.25 airy unit (AU) for Airyscan imaging and optimal resolution for Airyscan acquisition maintained (∼29.41 pixels/μm) regardless of frame size (17.8 – 73.2 μm, depending on experiment). An automatic Airyscan processing strength of 6.0 was applied to the image post-acquisition. Ten images were acquired per experiment and the experiment repeated three times.

### Nuclear localisation analysis

The nuclear:cytoplasmic mean GFP fluorescence intensity ratio was calculated from confocal images using the standard ImageJ toolkit. Nuclear fluorescence was calculated by masking the DAPI signal and measuring GFP fluorescence within the mask, both mean fluorescence intensity and total fluorescence. Mean cytoplasmic fluorescence intensity was calculated by subtracting total cellular GFP fluorescence by total nuclear GFP fluorescence and dividing the resulting value by the difference in area of the whole cell and nucleus.

### Co-localisation analysis

To quantify the co-localisation between GFP and subcellular markers (Calnexin, GM130, TGN46, EEA1 or LAMP1), Pearson’s correlation coefficient was calculated using the “Coloc 2” plugin in Fiji (ImageJ 1.51i) (83). First, images were split into GFP and marker channels and a region of interest (ROI) drawn around the cell using the GFP channel. Coloc2 was initiated using bisection threshold regression, a PSF of 3.0 pixels and a Costes randomisation value of 10. To analyse GFP overlap with the membrane marker FM4-64, line profiles were used. Ten-pixel wide, 5 μm-long line profiles were placed, with the membrane marker centred at 2.5 μm. Two to four line profiles were taken per cell and averaged. Fluorescence intensity was normalised to the maximum value for each cell. Ten cells were measured.

### Live cell imaging, fluorescence recovery after photobleaching (FRAP) and Förster resonance energy transfer (FRET) acquisition

10,000 cells were seeded per well into an 8-well Lab-Tek II chambered coverglass slide (Nunc) 48 h prior to imaging. In some experiments, FM4-64 (Thermo, 120 nM) or Hoechst 33342 (Thermo, 1 μg/ml) were applied immediately prior to imaging. FRAP acquisition was carried out using a Zeiss LSM 880 confocal microscope, with a Plan-Apochromat 63x oil immersion objective lens (NA = 1.4), controlled by ZEN2 software at 37 °C/5 % CO_2_. GFP was imaged using a 488 nm laser (1 – 5 % laser power), using bidirectional scanning at maximum scan speed and a 1 AU pinhole.

To monitor FRAP in the cytoplasm, a single cell was imaged at 10x zoom factor with a 256 × 256-pixel frame size. A 1 μm-wide ROI was photobleached with 40 iterations of a 488 nm laser (100 % laser power) and images acquired every 250 ms for 37.5 s. Alternatively, for a higher temporal resolution, a 64 × 64-pixel frame was used, and images acquired every 12.8 ms. Ten cells were imaged per experiment and three repeats performed.

To monitor FRAP in the nucleus, cells were imaged at 2.0x zoom factor with a 512 × 512-pixel frame size. A ROI was manually drawn around the nucleus (stained by Hoechst 33342) and photobleached with 40 iterations of the 488 nm laser (100 % laser power). Images were taken every 5 s until 10 successive images without an increase in nuclear fluorescence were acquired (typically 5 – 10 min). Three to four cells were imaged per experiment and the experiment repeated three times.

For FRET, cells were imaged at 2.0 – 4.0x zoom factor using a 512 × 512-pixel frame size. FM4-64 was bleached using 100 iterations of the 561 nm laser (100 % laser power) at the plasma membrane or within internal vesicles. Images were acquired every 0.6 s for 25 s.

### FRAP analysis

Analysis for FRAP data was adapted from (84). Images were exported to ImageJ for data acquisition using the FRAP Norm plugin. Three regions were plotted on the image: the photobleached ROI, delineated as the full width at half maximum of the encompassing cytoplasmic GFP fluorescence or the entire photobleached nucleus, for cytoplasmic and nuclear photobleaching experiments, respectively. A control region, placed elsewhere in the cell, was used to calculate the rate of photobleaching at each time point across the time series, relative to the maximal fluorescence intensity at *t* = 0. Lastly, a background region was placed outside the cell, which was subtracted from the other two regions at each time point. Therefore, at each time point, the ROI could be normalised to the photobleaching rate. Finally, fluorescence intensity within the ROI was normalised to pre-bleach fluorescence intensity to obtain the final recovery curve. Two parameters were derived from these recovery curves to quantify mobility. The mobile fraction, which defines the proportion of fluorescent protein that is mobile relative to the whole population of fluorescent protein initially in the ROI, was calculated using:

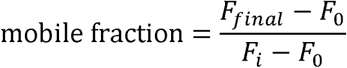

Where F_final_ = final fluorescence measurement, F_0_ = first post-bleach fluorescence measurement, and F_i_ = pre-bleach fluorescence measurement. The half-time describes the time taken for half maximal fluorescence recovery and was derived from a single exponential curve fitted to post-bleach measurements.

### FRET analysis

To analyse FRET, images were exported to ImageJ and analysed using the FRAP Norm plugin. GFP and FM4-64 fluorescence intensities were monitored within the photobleaching ROI for the duration of the time series and normalised against *t* = 0. FM4-64 signal was monitored to ensure photobleaching occurred. GFP fluorescence intensity before and after photobleaching was then statistically compared.

### Whole cell patch clamp recording

Whole cell patch clamp recordings were performed and analysed as described previously (85). Data were collected at a sampling rate of 50 kHz and filtered at 10 kHz. Linear leak currents were removed using P/6 subtraction (86). Series resistance was compensated by 40 %. Extracellular recording solution (physiological saline solution; PSS) contained (in mM): 144 NaCl, 5.4 KCl, 1 MgCl_2_, 2.5 CaCl_2_, 5 HEPES, 5.6 D-glucose, adjusted to pH 7.2 with KOH. For experiments involving cells pre-treated with drugs, the drug or vehicle was included in the PSS. The intracellular patch pipette solution contained (in mM): 5 NaCl, 145 CsCl, 2 MgCl_2_, 1 CaCl_2_, 10 HEPES and 11 EGTA, adjusted to pH 7.4 using CsOH. To assess current-voltage (I-V) relationships and activation-voltage relationships, cells were pre-pulsed at -120 mV for 250 ms before 5 mV depolarising steps for 50 ms in the range -80 mV to +30 mV. Steady-state inactivation was assessed at -10 mV for 50 ms following conditioning pre-pulse steps for 250 ms in the range -120 mV to -10 mV. Recovery from inactivation was assessed following depolarisation to 0 mV for 25 ms, holding at -120 mV for *t* ms, then second depolarisation at 0 mV for 25 ms. *t*: 1, 2, 3, 5, 7, 10, 15, 20, 30, 40, 50, 70, 100, 150, 200, 250, 350, 500 ms. In experiments involving TTX perfusion, TTX solution (1 μM in PSS) was exchanged for PSS (and vice versa) by performing three bath changes, with voltage clamp protocols run in each condition.

### RNA extraction and RT-qPCR

Total RNA was extracted from 35 mm dishes of confluent cells using RNeasy Mini kit (Qiagen), according to the manufacturer’s instructions. cDNA was generated from 1 µg of RNA using Reverse Transcriptase SuperScript III (RT SS III), random primers (Invitrogen), and dNTPs (Invitrogen). RNA, random primers, and dNTPs were incubated at 65°C for 5 min. Salt buffers, 0.1M DTT, RNase Out, and RT SS III were added and incubated at 25°C for 5 min, 50°C for 60 min, and at 70°C for 15 min. cDNA was either kept undiluted (*SCN1A, SCN4A, SCN8A*) or diluted 1:3 (*SCN2A, SCN3A, SCN9A*) in RNase-free water. Quantitative PCR was performed using SYBR Green (Applied Biosystems) and gene-specific primers (Integrated DNA Technologies) on a QuantStudio 7 Flex Real-Time PCR System (Applied Biosystems). Using the comparative threshold (2^-^ΔΔ^Ct^) method of quantification, gene-specific measurements of each cDNA sample were run in triplicate and compared to endogenous control gene *GAPDH*. Relative expression levels of genes were then normalized to the control condition (GFP) to determine gene expression.

### Data analysis

GraphPad Prism 8.0 was used for all curve fitting and statistical analyses. Normality of data was initially assessed using a D’Agostino and Pearson test. For normally distributed data, an unpaired Student’s t-test was used for pairwise comparisons and a one-way ANOVA with Tukey’s or Dunnett’s post hoc test used for multiple comparisons and data presented as mean ± SD. For data not following a normal distribution, pairwise comparisons were performed using a Mann-Whitney U-test and multiple comparisons were performed using a Kruskal-Wallis test with Dunn’s multiple comparison post hoc test. Results were considered significant if P < 0.05.

### Data availability

The datasets used and/or analysed during the current study are available from the corresponding author on reasonable request.

## Acknowledgements

The authors thank Michaela Nelson for technical assistance with establishing the cellular assays and the University of York Bioscience Technology Facility for their support with the confocal microscopy.

## Author contributions

AH, CB and WB contributed to the conception and design of the work. ASH, SLH, ALC, LLI, CGB and WJB contributed to acquisition, analysis, and interpretation of data for the work. ASH, SLH, ALC, LLI, CGB and WJB contributed to drafting the work and revising it critically for important intellectual content. All authors approved the final version of the manuscript.

